# Direct inference of base-pairing probabilities with neural networks improves RNA secondary structure prediction with pseudoknots

**DOI:** 10.1101/303172

**Authors:** Manato Akiyama, Yasubumi Sakakibara, Kengo Sato

**Affiliations:** Department of Biosciences and Informatics, Keio University, 3–14–1 Hiyoshi, Kohoku-ku, Yokoama 223–8522, Japan.

## Abstract

**Motivation:** Existing approaches for predicting RNA secondary structures depend on howto decompose a secondary structure into substructures, so-called the *architecture*, to define their parameter space. However, the architecture has not been sufficiently investigated especially for pseudoknotted secondary structures.

**Results:** In this paper, we propose a novel algorithm to directly infer base-pairing probabilities with neural networks that does not depend on the architecture of RNA secondary structures, followed by performing the maximum expected accuracy (MEA) based decoding algorithms; Nussinov-style decoding for pseudoknot-free structures, and IPknot-style decoding for pseudoknotted structures. To train the neural networks connected to each base-pair, we adopt a max-margin framework, called structured support vector machines (SSVM), as the output layer. Our benchmarks for predicting RNA secondary structures with and without pseudoknots show that our algorithm achieves the best prediction accuracy compared with existing methods.

**Availability:** The source code is available at https://github.com/keio-bioinformatics/neuralfold/.

## Introduction

Recent studies have unveiled that functional non-coding RNAs (ncRNAs) play essential roles such as transcriptional regulation and guiding modification, resulting in various biological processes ranging from development and cell differentiation to the cause of diseases (Hirose *et al.*, 2014). Since it is well-known that functions of ncRNAs are deeply related to their structures rather than primary sequences, discovering the structures of ncRNAs leads to understanding the functions of ncRNAs. However, there are severe difficulties in experimental assays to determine RNA tertiary structures such as nuclear magnetic resonance (NMR) and X-ray crystal structure analysis due to high experimental costs and size limits of measurements on RNA. Therefore, instead of such experimental assays, we frequently perform computational prediction of RNA secondary structures, which is defined as a set of base-pairs consisting of hydrogen bonds between nucleotides.

The most popular approach for predicting RNA secondary structures is based on thermodynamic models such as Turner’s nearest neighbor model (Schroeder and Turner, 2009; Turner and Mathews, 2010), which defines characteristic substructures such as hairpin loops and base-pair stacking. Free energy of each substructure has been determined by experimental methods such as the optical melting experiment (Schroeder and Turner, 2009). The free energy of an RNA secondary structure is calculated by summing up the free energy of substructures into which it is decomposed. We can employ the dynamic programming technique to find the optimal secondary structure that minimizes the free energy for a given RNA sequence. A number of tools including UNAfold (Zuker, 1989), RNAfold (Lorenz *et al.*, 2011) and RNAstructure (Reuter and Mathews, 2010) have adopted this approach.

An alternative approach utilizes machine learning techniques, which train scoring parameters for decomposed substructures from reference structures, instead of the experimental techniques. This approach has successfully been adopted by CONTRAfold (Do *et al.*, 2006, 2007), Simfold (Andronescu *et al.*, 2007, 2010a), ContextFold (Zakov *et al.*, 2011) and so on, and thus has enabled us to predict more accurate RNA secondary structures.

Another important aspect of the RNA secondary structure prediction is the choice of the decoding algorithm, which finds an optimal secondary structure from all the possible secondary structures. A classic decoding algorithm is the minimum free energy (MFE) based algorithm for the thermodynamic approach, or the maximum likelihood estimation (MLE) based algorithm for the machine learning based approach, which finds a secondary structure that minimizes (resp. maximizes) the free energy (resp. probability or scoring function). Another choice is the posterior decoding algorithm based on the maximum expected accuracy (MEA) principle, which is known to be one of the effective approaches for many high-dimensional combinatorial optimization problems (Carvalho and Lawrence, 2008). Since we usually evaluate prediction of RNA secondary structures by base-pair-wise accuracy measures, the MEA-based decoding algorithms utilize posterior base-pairing probabilities that can be calculated by McCaskill algorithm (McCaskill, 1990) or the inside-outside algorithmfor stochastic context-free grammars. CONTRAfold and CentroidFold (Hamada *et al.*, 2009; Sato *et al.*, 2009) have successfully implemented the MEA-based decoding algorithm for predicting RNA secondary structures.

Pseudoknot is one of the important structural elements in RNA secondary structures. A secondary structure includes a pseudoknot if at least two arcs (hydrogen bonds) drawn above the primary sequence cross each other (Fig. 1). Many RNAs such as rRNAs, tmRNAs and viral RNAs form pseudoknotted secondary structures (van Batenburg *et al.*, 2001). It is known that pseudoknots are involved in the regulation of translation and splicing, and ribosomal frame shifting (Staple and Butcher, 2005; Brierley *et al.*, 2007). Furthermore, pseudoknots assist folding into 3D structures in many cases (Fechter *et al.*, 2001). Therefore, pseudoknots cannot be ignored for structural and functional analysis of RNAs.

**Fig. 1.**
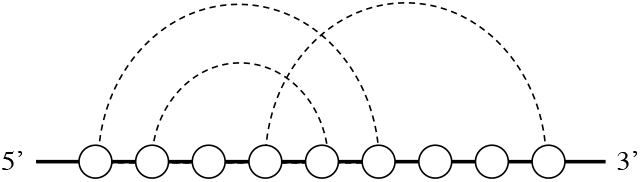
An example of pseudoknots.

However, all of the above-mentioned algorithms cannot consider pseudoknotted secondary structures due to computational complexity. It has been proven that the problem of finding the MFE structure including arbitrary pseudoknots is NP-hard (Akutsu, 2000; Lyngsø and Pedersen, 2000). Therefore, practically available algorithms for predicting pseudoknotted RNA secondary structures fall into one of the following two approaches; the exact algorithms for a limited class of pseudoknots such as PKNOTS (Rivas and Eddy, 1999), NUPACK (Dirks and Pierce, 2003, 2004) and pknotsRG (Reeder and Giegerich, 2004), and the heuristic algorithms that do not guarantee the optimal structure such as ILM (Ruan *et al.*, 2004), HotKnots (Andronescu *et al.*, 2010b; Ren *et al.*, 2005), FlexStem (Chen *et al.*, 2008) and ProbKnot (Bellaousov and Mathews, 2010).

We have previously developed IPknot, which enables us fast and accurate prediction of RNA secondary structures with pseudoknots using integer programming (Sato *et al.*, 2011). IPknot adopts the MEA-based decoding algorithm that utilizes base-pairing probabilities combined with an approximation of decomposing a pseudoknotted structure into hierarchical pseudoknot-free structures. Prediction performance of IPknot is sufficiently good in speed and accuracy as compared with the heuristic algorithms, and is much faster than the exact algorithms.

Both the thermodynamic approach and the machine learning based approach depend on how to decompose a secondary structure into substructures, so-called the *architecture* in (Rivas, 2013), to define their parameter space. The Turner’s nearest neighbor model is the most well-studied architecture for pseudoknot-free secondary structures, meanwhile the energy model for pseudoknotted secondary structures has not been sufficiently investigated except for the Dirks–Pierce model (Dirks and Pierce, 2003, 2004) and the Cao–Chen model (Cao and Chen, 2006) for limited classes of pseudoknots. To the best of our knowledge, an effective and efficient procedure to find a suitable architecture that can predict RNA secondary structures more accurately is still unknown.

In this paper, we propose a novel algorithm to directly infer base-pairing probabilities with neural networks instead of the McCaskill algorithm or the inside-outside algorithm, which depend on the architecture of RNA secondary structures. Then, we employ the inferred base-pairing probabilities as part of the MEA-based scoring function for the decoding algorithms; Nussinov-style decoding for pseudoknot-free structures and IPknot-style decoding for pseudoknotted structures. To train the neural networks connected to each base-pair, we adopt a max-margin framework, called structured support vector machines (SSVM), as the output layer. We implement two types of neural networks connected to each base-pair; bidirectional recursive neural networks (BiRNN) over tree structures and multilayer feedforward neural networks (FNN) with *k*-mer contexts around both of paired bases. Our benchmarks for predicting RNA secondary structures with and without pseudoknots show that our algorithm achieves the best prediction accuracy compared with existing methods.

The major advantages of our work are summarized as follows: (i) our algorithm enables us to accurately predict RNA secondary structures with and without pseudoknots, and (ii) our algorithm assumes no prior knowledge of the architecture that defines the decomposition of RNA secondary structures and thus the parameter space.

## 2 Methods

### 2.1 Preliminaries

Let Σ = {A, C, G, U} and Σ* denote the set of all finite RNA sequences consisting of bases in Σ. For a sequence *x* = *x*_1_*x*_2_ ⋯ *x_n_* ∈ Σ*, let |*x*| denote the number of bases appearing in *x*, which is called the length of *x*. Let 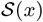 be a set of all possible secondary structures of *x*. A secondary structure 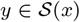 is represented as a |*x*| × |*x*| binary-valued triangular matrix *y* = (*y_ij_*)*i*<*j*, where *y_ij_* = 1 if and only if bases *x_i_* and *x_j_* form a base-pair composed by hydrogen bonds including the Watson-Crick base-pairs (A-U and G-C), the Wobble base-pairs (G-U).

### 2.2 MEA-based scoring function

We employ the maximum expected accuracy (MEA) based scoring function that has been originally used for IPknot (Sato *et al.*, 2011).

We assume that a secondary structure 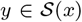 can be decomposed into a set of pseudoknot-free substructures (*y*^(1)^, *y*^(2)^, …, *y*^(*m*)^) that satisfies the following conditions: (1) 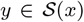 should be decomposed into a mutually-exclusive set, that is, for all 1 ≤ *i* < *j* ≤ |*x*|, 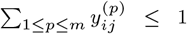; and (2) every base pair in *y*^(*p*)^ should be pseudoknotted to at least one base pair in *y*^(*q*)^ for ∀*q* < *p*. Each pseudoknot-free substructure *y*^(*p*)^ is said to belong to the *level p*. For any RNA secondary structure 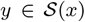, there exists a positive integer *m* such that *y* can be decomposed into *m* pseudoknot-free substructures. From this viewpoint, we can say that the above decomposition enables our method to model arbitrary pseudoknots.

First, we define a gain function of 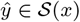 with regard to the correct secondary structure 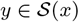 as follows:

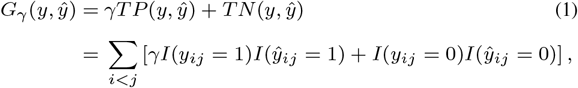

where *γ* > 0 is a weight parameter for base pairs, *TP* and *TN* denote the numbers of true positives (base pairs) and true negatives (non-base pairs), respectively, and *I*(*condition*) is the indicator function that takes a value of 1 or 0 depending on whether the *condition* is true or false.

Our objective is to find a secondary structure *ŷ* that maximizes the expectation of the gain function (1) under a given probability distribution over the space 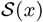 of pseudoknotted secondary structures:

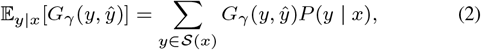

where *P*(*y* | *x*) is a probability distribution of RNA secondary structures including pseudoknots. It has been proven that the γ-centroid estimator (2) enables us to decode secondary structures accurately from a given probability distribution (Hamada *et al.*, 2009).

We approximate the expected gain function (2) by the sum of the expected gain functions for each level of pseudoknot-free substructures (*ŷ*^(1)^, …, *ŷ*^(*m*)^) in the decomposed set of a pseudoknotted structure 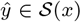, and thus simultaneously find a pseudoknotted structure *ŷ* and its decomposition (*ŷ*^(1)^, …, *ŷ*^(*m*)^) that maximize:

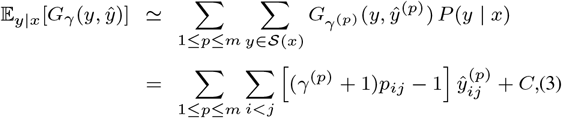

where *γ*^(*p*)^ > 0 is a weight parameter for base pairs at the level *p*, and *C* is a constant independent of *ŷ* (see the Supplementary Material of (Hamada *et al.*, 2009) for the derivation). The base-pairing probability *p_ij_* is the probability that the base *x*_*i*_ is paired with *x_j_*. As seen in Sec. 2.4, we employ one of the three algorithms for calculating base-pairing probabilities.

It is worth mentioning that IPknot can be regarded as an extension of CentroidFold (Hamada *et al.*, 2009). If we let the number of decomposed levels *m* = 1, the approximate expected gain function (3) is identical to the γ-centroid estimator used in CentroidFold.

### 2.3 Decoding algorithms

#### 2.3.1 Nussinov-style decoding algorithm for pseudoknot-free structures

For pseudoknot-free secondary structure prediction, we find *ŷ* that maximizes the expected gain (3) with *m* = 1 under the constraints on base-pairs, that is,

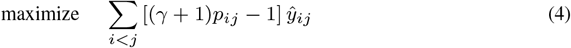

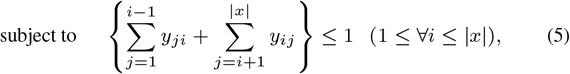

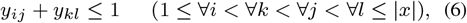

This integer programming problem (IP) can be solved by using the following dynamic programming similar to Nussinov algorithm (Nussinov *et al.*, 1978):

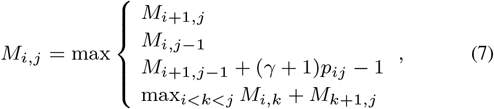

and tracing back from *M*_1,|*x*|_.

#### 2.3.2 IPknot-style decoding algorithm for pseudoknotted structures

Maximization of the approximate expected gain (3) can be solved by the IP problem as follows:

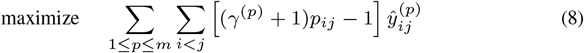

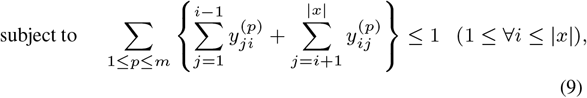

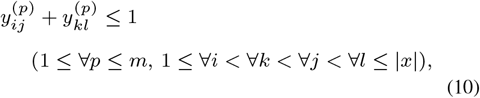

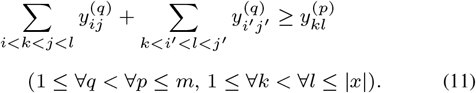

Note that due to Eq. (3), we need to consider only base pairs 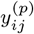 whose base-pairing probabilities *p_ij_* are larger than *θ*^(*p*)^ = 1/(*γ*^(*p*)^ + 1). The constraint (9) means that each base *x_i_* can be paired with at most one base. The constraint (10) disallows pseudoknots within the same level *p*. The constraint (11) ensures that each base pair at the level *p* is pseudoknotted to at least one base pair at every lower level *q* < *p*. We set *m* = 2 by default according to IPknot’s default. This suggests that the predicted structure can be decomposed into two pseudoknot-free secondary structures.

### 2.4 Inferring base-paring probabilities

Our scoring function (3) described in Sec. 2.2 is calculated by using base-pairing probabilities *p_ij_*. In this section, we introduce two approaches for computing base-pairing probabilities. The first approach is a traditional one that is based on the probability distribution of RNA secondary structures, e.g., the McCaskill model (McCaskill, 1990) for pseudoknot-free structures and its extension to pseudoknotted structures such as the Dirks–Pierce model (Dirks and Pierce, 2003, 2004). The second approach proposed in this paper directly calculates base-pairing probabilities using neural networks.

#### 2.4.1 Traditional models for base-pairing probabilities

The base-pairing probability *p_ij_* is defined as:

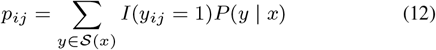

from a probability distribution *P*(*y* | *x*) over a set 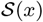 of secondary structures *with* or *without* pseudoknots.

For predicting pseudoknot-free structures, the McCaskill model (McCaskill, 1990) can be mostly used as *P*(*y* | *x*) combined with the Nussinov-style decoding algorithm described in Sec. 2.3.1. The computational complexity of calculating Eq. (12) for the McCaskill model is *O*(|*x*|^3^) for time and *O*(|*x*|^2^) for space by using the dynamic programming technique. This model has been implemented previously as CentroidFold (Hamada *et al*, 2009; Sato *et al*, 2009).

For predicting pseudoknotted structures, we can select *P*(*y* | *x*) from several models. A naïve model is the use of the probability distribution *with* pseudoknots as well as Eq. (2) in spite of high computational costs, e.g., the Dirks–Pierce model (Dirks and Pierce, 2003, 2004) for a limited class of pseudoknots, whose computational complexity is *O*(|*x*|^5^) for time and *O*(|*x*|^4^) for space. Alternatively, we can employ a probability distribution *without* pseudoknots for each decomposed pseudoknot-free structure such as the McCaskill model. Furthermore, to boost the prediction accuracy, we can utilize a heuristic algorithm, the iterative refinement, that refines the base-pairing probability matrix from the distribution without pseudoknots. See (Sato *et al.*, 2011) for more details. These three models have been implemented as IPknot (Sato *et al.*, 2011).

#### 2.4.2 Neural network models

We propose two neural network architectures for calculating base-pairing probabilities instead of the probability distribution over RNA secondary structures.

The first architecture is the bidirectional recursive neural network (BiRNN) over tree structures as shown in Fig. 2. The BiRNN consists of the three matrices: (a) the inside RNN matrix, (b) the outside RNN matrix and (c) the inside-outside matrix for outputting base-pairing probabilities, each of whose element contains a network layer (indicated by a circle) with 80 hidden nodes. Each layer in the inside (resp. outside) matrix is recursively calculated from connected source layers as like the inside (resp. outside) algorithm for the stochastic context-free grammars (SCFG). The ReLU activation function is applied before input to each recursive node. The base-pairing probability at each position is calculated from the corresponding layers in the inside and outside matrices with the sigmoid activation function. Our implementation of BiRNN assumes a simple RNA grammar

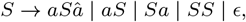

where *a* ∈ Σ, *a* and *â* stand for paired bases, *S* is the start non-terminal symbol, *ϵ* is the empty string.

**Fig. 2.**
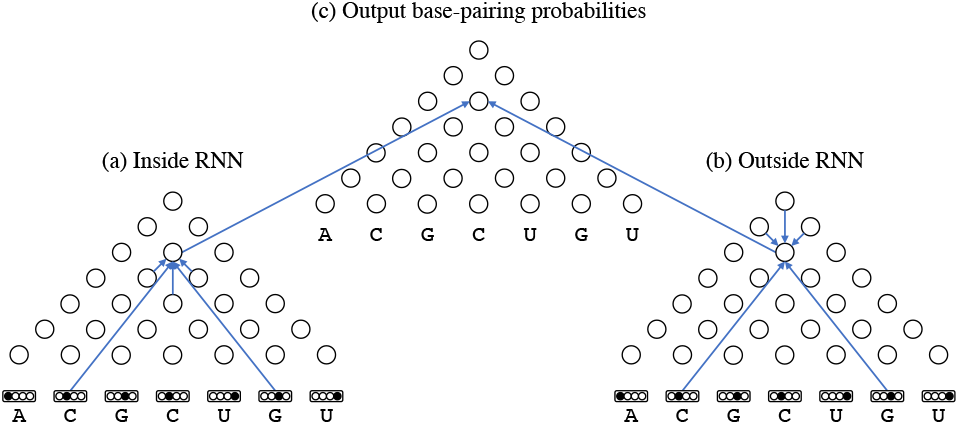
A bidirectional recursive neural network for calculating base-pairing probabilities. Arrows indicate the network parameters of neural networks.

The second architecture employs simple multilayer feedforward neural networks (FNN). To calculate the base-pairing probability *p_ij_*, an FNN inputs two *k*-mers around *i*-th and *j*-th bases as shown in Fig. 3. Each base is encoded by the one-hot encoding of nucleotides and an additional node that indicates the end of the loop, which should be active for *x_l_* s.t. *l* ≥ *j* in the left *k*-mer around *x*_*i*_ or *x*_*l*_ s.t. *l* ≤ *i* in the right *k*-mer around *x_j_*. We can expect that this encoding embeds the length of loops and the contexts around the openings and closings of helices. We set *k* = 81 for the *k*-mer context length default (See for more details in Sec. 3.4). We construct two hidden layers consisting of 200 and 50 nodes with the ReLU activation function, and one output node with the sigmoid activation function to output base-pairing probabilities.

**Fig. 3.**
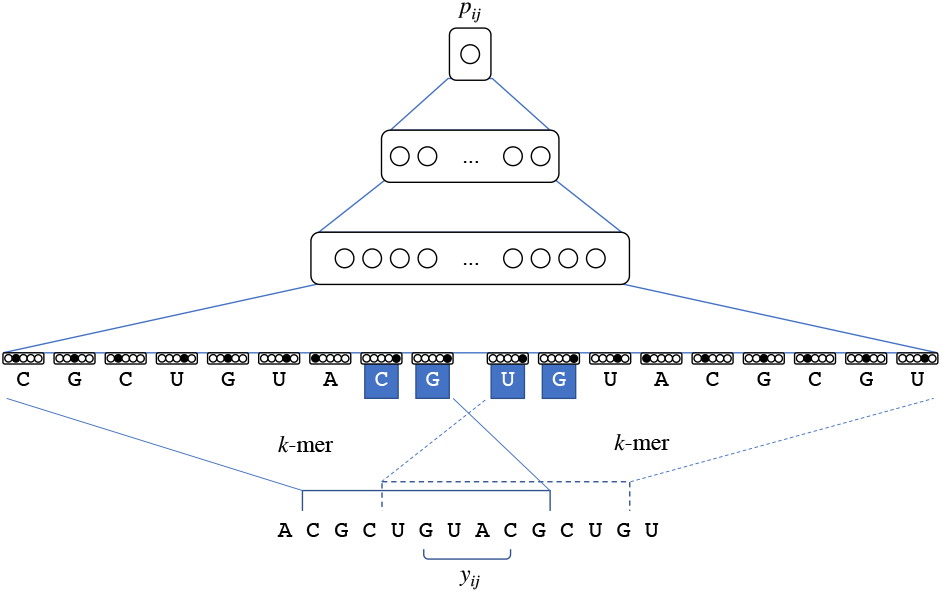
A feedforward neural network with *k*(= 9)-mer contexts around *x_i_* and *x_j_* calculates the base-pairing probability *p_ij_*. The end-of-loop nodes of the highlighted nucleotides are activated since they are beyond the paired bases.

Note that the FNN model depends on no assumption of RNA secondary structures, while the BiRNN model assumes an RNA grammar that considers no pseudoknots. Instead, the FNN model can take longer contexts around each base-pair into consideration by using longer *k*-mers.

### 2.5 Learning algorithm

To optimize the network parameters *λ*, we employ a max-margin framework called structured support vector machines (SSVM) (Tsochantaridis *et al.*, 2005). Given a training dataset 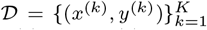, where *x*^(*k*)^ is the *k*-th RNA sequence and 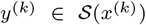 is the correct secondary structure for the *k*-th sequence *x*^(*k*)^, we aim to find λ that minimizes the objective function

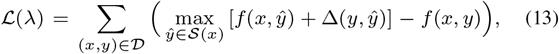

where *f*(*x, y*) is the scoring function of RNA secondary structure 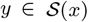 for a given RNA sequence *x* ∈ Σ*, that is, Eq. (4) for the Nussinov-style decoding, or Eq. (8) for the IPknot-style decoding. Flere, Δ(*y, ŷ*) is a loss function of *ŷ* for *y* defined as

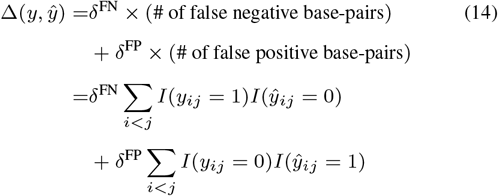

where *δ*^FN^ and *δ*^FP^ are tunable hyperparameters to control the trade-off between sensitivity and specificity for learning the parameters. We used *δ*^FN^ = *δ*^FP^ = 0.1 by default. In this case, we can calculate the first term of Eq. (13) using the Nussinov-style decoding algorithm or the IPknot-style decoding algorithm modified by the loss-augmented inference (Tsochantaridis *et al.*, 2005).

To minimize the objective function (13), we can apply stochastic subgradient descent (Fig. 4) or its variant. We can calculate the gradients with regard to the network parameters λ for the objective function (13) using the gradients with regard to *p_ij_* by the chain rule of differential. This means that the prediction errors occurred by the decoding algorithm backpropagate to the neural network that calculates base pairing probabilities through the connected base pairs.

**Fig. 4.**
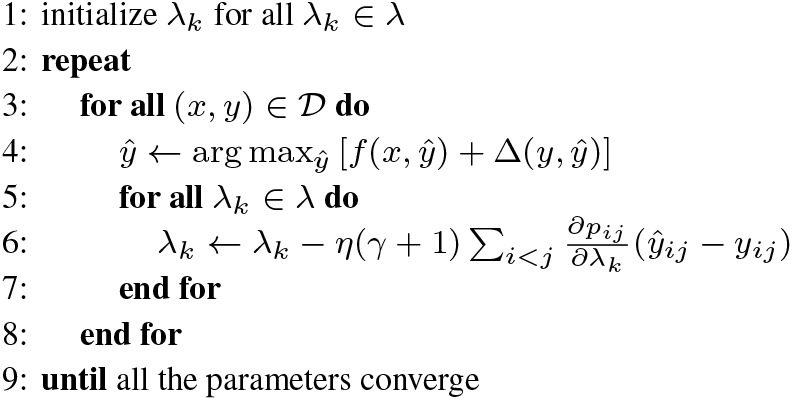
The stochastic subgradient descent algorithm for SSVMs. *η* > 0 is the predefined learning rate.

## 3 Results

### 3.1 Implementation

Our algorithm was implemented as a program called Neuralfold, which is short for the Neural network based RNA FOLDing algorithm. We employ CPLEX IP solver^1^ to solve the IP problem (8)–(11). The source code is available at https://github.com/keio-bioinformatics/neuralfold/.

### 3.2 Datasets

We evaluated our algorithm with the Nussinov-style decoding algorithm for predicting pseudoknot-free RNA secondary structures on two datasets: TrainSetB and TestSetB assembled from Rfam (Gardner *et al.*, 2011), which contain 22 families with 3D structures (Rivas, 2013). TrainSetB and TestSetB include sequences from Rfam seed alignments with no more than 70% identity among each other. TestSetB is made up of 22 RNA families and its composition is 14 5.8SrRNAs, 18 U1 spliceosomal RNAs, 45 U4 spliceosomal RNAs, 233 riboswitches (from seven different families), 116 cis regulatory elements (from nine different families), three ribozymes, and one bacteriophage pRNA. TestSetB contains 430 sequences. There are 52,097 residues in all, of which 22,728 bases (43.6%) form base pairs. The sequence length is in the range of 27 to 244 nt and the average length is 121 nt. TestSetB contains 8.3% noncanonical base pairs. TrainSetB also consists of 22 RNA families as same as TestSetB, by selecting the sequences dissimilar with TestSetB. TrainSetB contains 1094 sequences. There are 112,398 residues in all, of which 52,065 bases (46.3%) form base pairs. The sequence length is in the range of 27 to 237 nt and the average length is 103 nt. TrainSetB contains 4.3% noncanonical base pairs.

We also evaluated our algorithm with the IPknot-style decoding algorithm for predicting pseudoknotted RNA secondary structures on pk168 dataset (Huang and Ali, 2007), which was compiled from PseudoBase (van Batenburg *et al.*, 2001), This dataset includes 16 categories of 168 pseudoknotted sequences whose lengths are <140 nt.

### 3.3 Prediction performance

We evaluated the accuracy of predicting RNA secondary structures through the sensitivity (SEN) and the positive predictive value (PPV), defined as:

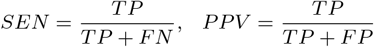

where *TP* is the number of correctly predicted base-pairs (true positives), *FP* is the number of incorrectly predicted basepairs (false positives), and *FN* is the number of base-pairs in the true structure that were not predicted (false negatives). We also used the F-value as the balanced measure between SEN and PPV, which is defined as their harmonic mean:

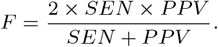

We conducted computational experiments on the datasets described in the previous section using the Nussinov-style decoding algorithm with the McCaskill model and the neural network models: the BiRNN model and the FNN model. We employed CentroidFold as the Nussinov decoding algorithm with the McCaksill model. We performed experiments on TestSetB using the parameters trained from TrainSetB. As shown in Table 1, the neural network models achieved better accuracy compared with the traditional model. Hereafter, we adopt the FNN model with fc-mer contexts as the default model of Neuralfold.

**Table 1.** The accuracy on the pseudoknot-free datasets.

| Implementation | Model | SEN | PPV | F |
| --- | --- | --- | --- | --- |
| Neuralfold | BiRNN | 0.649 | 0.601 | 0.624 |
| Neuralfold | FNN | 0.600 | 0.700 | 0.646 |
| CentroidFold | McCaskill | 0.513 | 0.544 | 0.528 |

The other computational experiments on the pk168 pseudoknotted dataset were conducted using the IPknot-style decoding algorithm with the McCaskill model with and without the iterative refinement, and the Dirks–Pierce model as well as Neuralfold with the FNN model. Table 2 shows that the FNN model is comparable to IPknot with the Dirks–Pierce model for pseudoknots, and better than the McCaskill model with and without the iterative refinement.

**Table 2.** The accuracy on the pseudoknotted datasets.

| Implementation | Model | SEN | PPV | F |
| --- | --- | --- | --- | --- |
| Neuralfold | FNN | 0.801 | 0.762 | 0.781 |
| IPknot | McCaskill w/o refine. | 0.619 | 0.710 | 0.661 |
| IPknot | McCaskill w/ refine. | 0.753 | 0.684 | 0.717 |
| IPknot | Dirks–Pierce | 0.809 | 0.749 | 0.778 |

refine.: the iterative refinement

Table 3 shows the computation time for various lengths of sequences; PKB229 and PKB134inthepk168 dataset, and ASE_00193, CRW_00614 and CRW_00774 in RNA STRAND database (Andronescu *et al.*, 2008). This shows that the computation time for predicting pseudoknotted secondary structure of the FNN model is comparably fast to IPknot with the Dirks–Pierce model.

**Table 3.** Computation time for calculating base-pairing probabilities of various lengths of sequences.

| ID | PKB229 | PKB134 | ASE_00193 | CRW_00614 | CRW_00774 |
| --- | --- | --- | --- | --- | --- |
| length (nt) | 67 | 137 | 301 | 494 | 989 |
| Neuralfold (FNN) | 3.30s | 27.78s | 44.73s | 60.22s | 3m4.2s |
| IPknot (w/o refine.) | 0.01s | 0.05s | 0.18s | 0.55s | 2.64s |
| (w/ refine.) | 0.03s | 0.08s | 0.31s | 1.03s | 5.86s |
| (D&P) | 8.36s | 9m4.7s | n/a | n/a | n/a |

Computation time was measured on Linux OS v2.6.32 with Intel Xeon E5-2680 (2.80 GHz) and 64 GB memory.

### 3.4 Effects of context length

We evaluated the prediction accuracy of the FNN model on the pseudoknot-free dataset and the pk168 dataset for several lengths of *k*-mers to be input to neural networks. Figures 5 and 6 show the accuracy for each feature representation with different *k*-mer lengths *k* = {3, 7, 11, 15, 19, 21, 41, 61, 81, 101, 121}. This indicates that the accuracy is improved mostly when the length of the *k*-mer is 81, and the difference of the accuracy on *L* ≥ 81 is negligible.

**Fig. 5.**
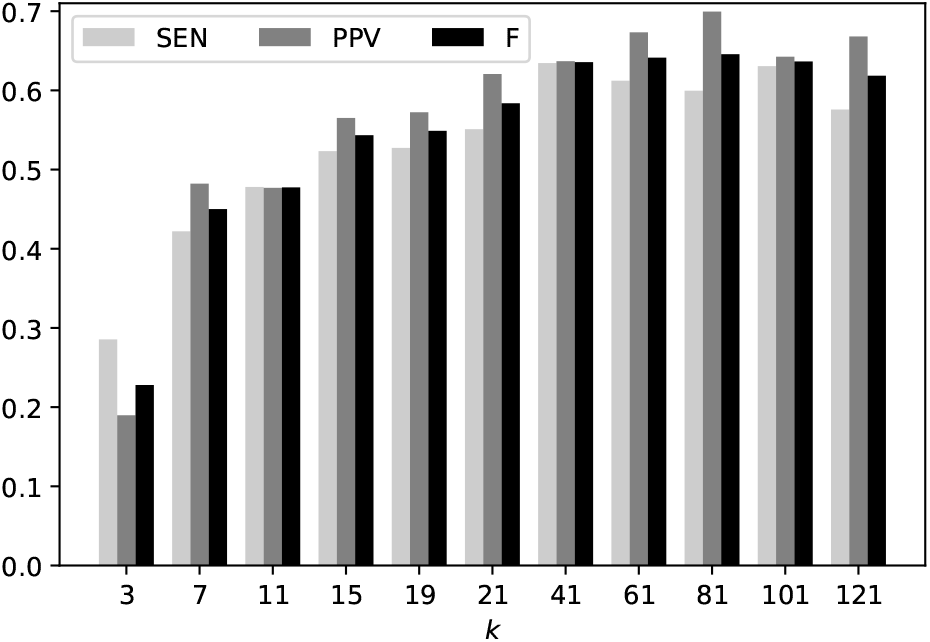
The accuracy of the FNN model with different lengths of *k*-mers on the pseudoknot-free dataset.

**Fig. 6.**
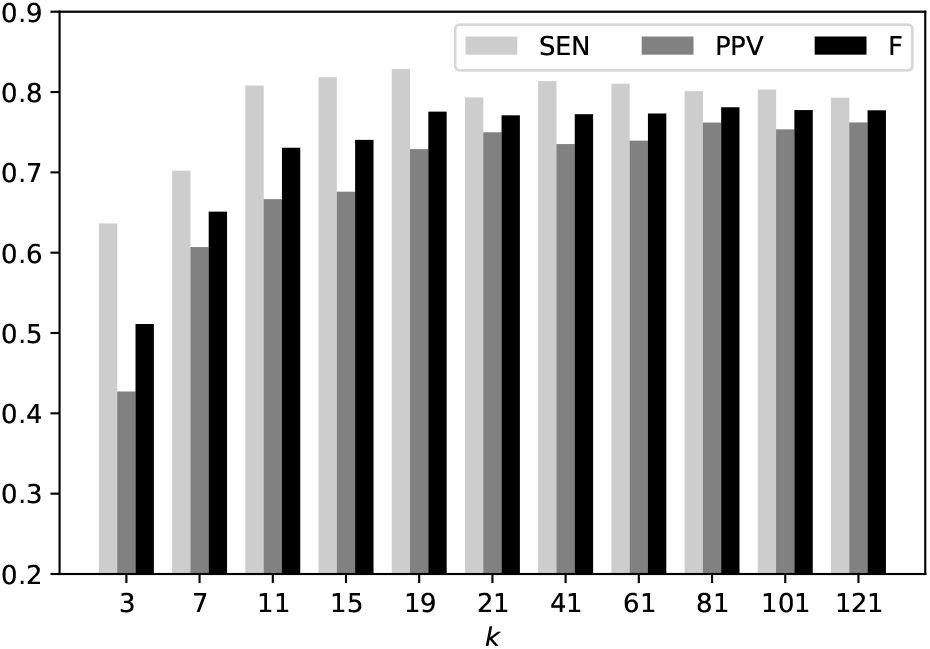
The accuracy of the FNN model with different lengths of *k*-mers on the pseudoknotted dataset.

### 3.5 Comparison with competitive methods for predicting pseudoknot-free secondary structures

We compared our algorithm with the other competitive methods for predicting pseudoknot-free RNA secondary structures including CentroidFold (Hamada *et al.*, 2009; Sato *et al.*, 2009), CONTRAfold (Do *et al.*, 2006, 2007), RNAfold in the Vienna RNA package (Lorenz *et al.*, 2011) and ContextFold (Reeder and Giegerich, 2004). For the posterior decoding methods with the trade-off parameter *γ* in Eq. (4), we used *γ* ∈ {2^*n*^ | *n* ∈ ℤ, –5 ≤ *n* ≤ 10}. Figure 7 shows PPV-SEN plots for each method, indicating that our algorithm works accurately on the pseudoknot-free dataset.

**Fig. 7.**
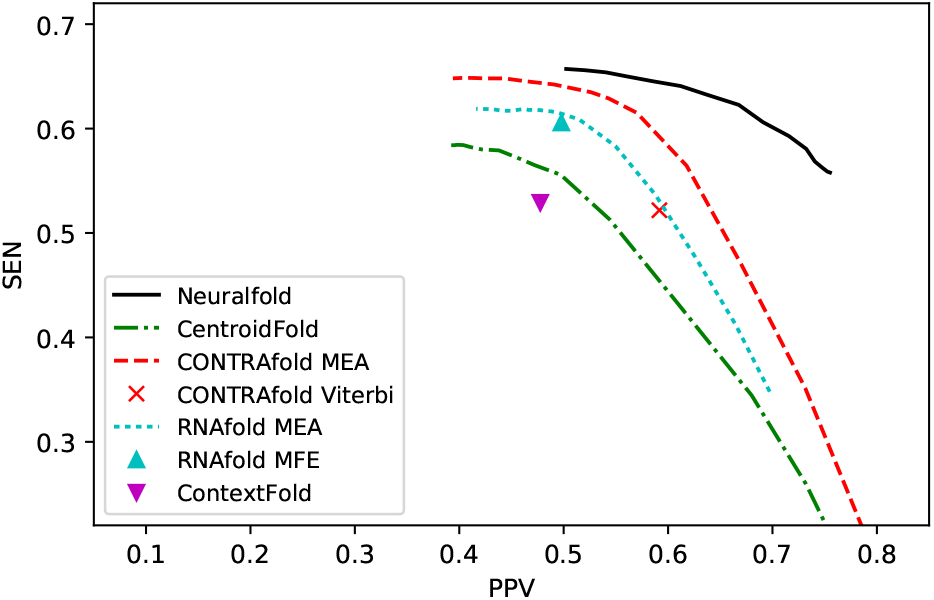
PPV-SEN plots comparing our algorithm with the competitive methods on the pseudoknot-free dataset.

### 3.6 Comparison with competitive methods for predicting pseudoknotted secondary structures

We also compared our algorithm with the other competitive methods for predicting pseudoknotted secondary structures including IPknot (Sato *et al.*, 2011), ProbKnot (Bellaousov and Mathews, 2010), FlexStem (Chen *et al.*, 2008), HotKnots (Andronescu *et al.*, 2010b; Ren *et al.*, 2005), pknotsRG (Reeder and Giegerich, 2004), ILM (Ruan *et al.*, 2004), NUPACK (Dirks and Pierce, 2003, 2004) and PKNOTS (Rivas and Eddy, 1999) as well as the methods for predicting pseudoknot-free secondary structures including CentroidFold and RNAfold. Neuralfold performed 10-fold cross validation on the pk168 dataset. Figure 8 shows PPV-SEN plots for each method, indicating that our algorithm works accurately on the pk168 dataset.

**Fig. 8.**
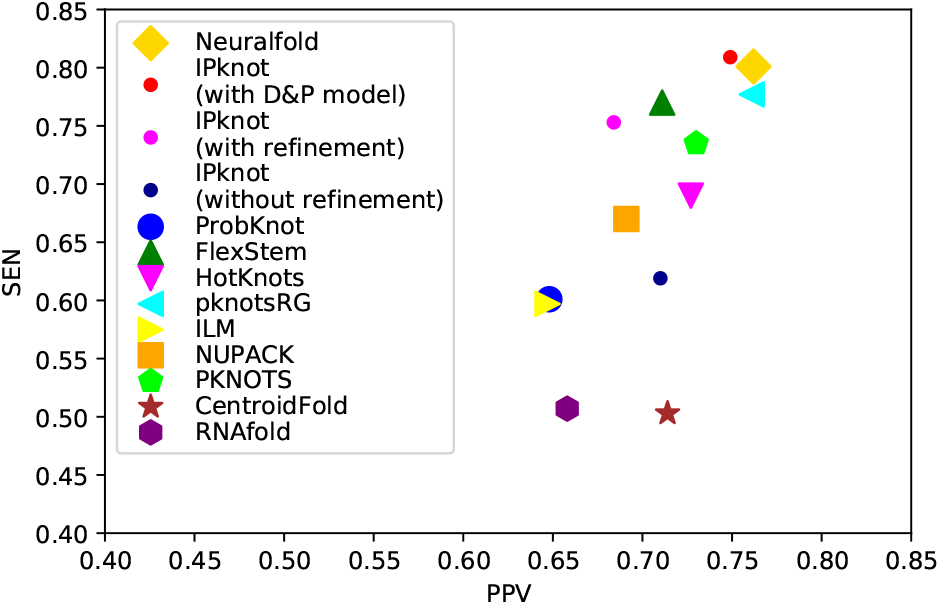
PPV-SEN plots comparing our algorithm with the competitive methods on the pseudoknotted dataset. We set γ^(1)^ =2, γ^(2)^ = 2 for Neuralfold, γ^(1)^ =2, γ^(2)^ = 4 for IPknot with D&P model, γ^(1)^ =2, γ^(2)^ = 16 for IPknot with/without refinement, and γ = 2 for CentroidFold.

## 4 Discussion

We propose a novel algorithm for directly inferring base-pairing probabilities with neural networks, which enables us to predict RNA secondary structures accurately. Sato *et al.* (2011) have previously proposed the iterative refinement algorithm for base-pairing probabilities, which refines the base-pairing probabilities calculated by the McCaskill algorithm so as to fit for pseudoknotted secondary structure prediction. The direct inference of base-pairing probabilities with neural networks is a similar approach to the iterative refinement algorithm in the sense that both directly update base-pairing probabilities, followed by the IPknot-style decoding algorithm using the base-pairing probabilities. Although the iterative refinement algorithm could fortunately improve the prediction accuracy of IPknot partly, it should be stated that the iterative refinement algorithm is an ad-hoc algorithm since there exists no theoretical guarantee. Meanwhile, the neural networks that infer base-pairing probabilities are trained from given reference secondary structures by the max-margin framework, meaning that we can theoretically expect that the neural network models improves the secondary structure prediction. In fact, Table 2 shows that our algorithm achieved not only better accuracy than the iterative refinement algorithm, but is also comparable to that of the Dirks–Pierce model, which can calculate exact base-pairing probabilities for a limited class of pseudoknots.

The direct inference of base-pairing probabilities with neural networks presented in this paper is the first algorithm that can be trained for pseudoknotted secondary structures except for HotKnots 2.0 (Andronescu *et al.*, 2010b), which finds a pseudoknotted secondary structure by an MFE-based heuristic decoding algorithm with energy parameters of the Dirks–Pierce model or the Cao–Chen model trained from pseudoknotted reference structures. One of the advantages of our algorithm over HotKnots 2.0 is that no assumption on the architecture of RNA secondary structures is required. In other words, our model can be trained from arbitrary classes of pseudoknots, while HotKnots cannot be trained from more complicated classes of pseudoknots than the one that the model had assumed. Furthermore, our algorithm can compute base-pairing probabilities, which can be applicable for various applications of RNA informatics such as family classification (Sato *et al.*, 2008; Morita *et al.*, 2009), RNA-RNA interaction prediction (Kato *et al.*, 2010) and simultaneous aligning and folding (Sato *et al.*, 2012). Accurate base-pairing probabilities calculated by our algorithm can improve the quality of such applications.

The FNN model takes two *k*-mers around each base-pair as input to infer its base-pairing probability, where *k* is the context length to model the length of loops and the contexts around the openings and closings of helices. Here, we can see in Figure 9 how different the context *k*-mer lengths will affect the prediction of pseudoknotted secondary structure. Consider the input bases when calculating the base pairing probability of the blue-highlighted base pair (AU) using the FNN model. The FNN model with the context length *k*=11 takes 5 bases on both the upstream and the downstream of the base *i* and *j* as input. As seen in Figure 9 (bottom), the distances from the bases A and U are 10 and 13 to the stem 2, respectively. This means that all the bases of the stem 2 are NOT completely located within the context length *k*=11 around the base pair AU. On the other hand, for the FNN model with the context length *k*=41, all the bases of the stem 2 are completely located within the context around the base pair AU. This leads the FNN model to correctly predict the base pair AU, suggesting that longer context length enables to consider dependency between stems in pseudoknotted substructures.

**Fig. 9.**
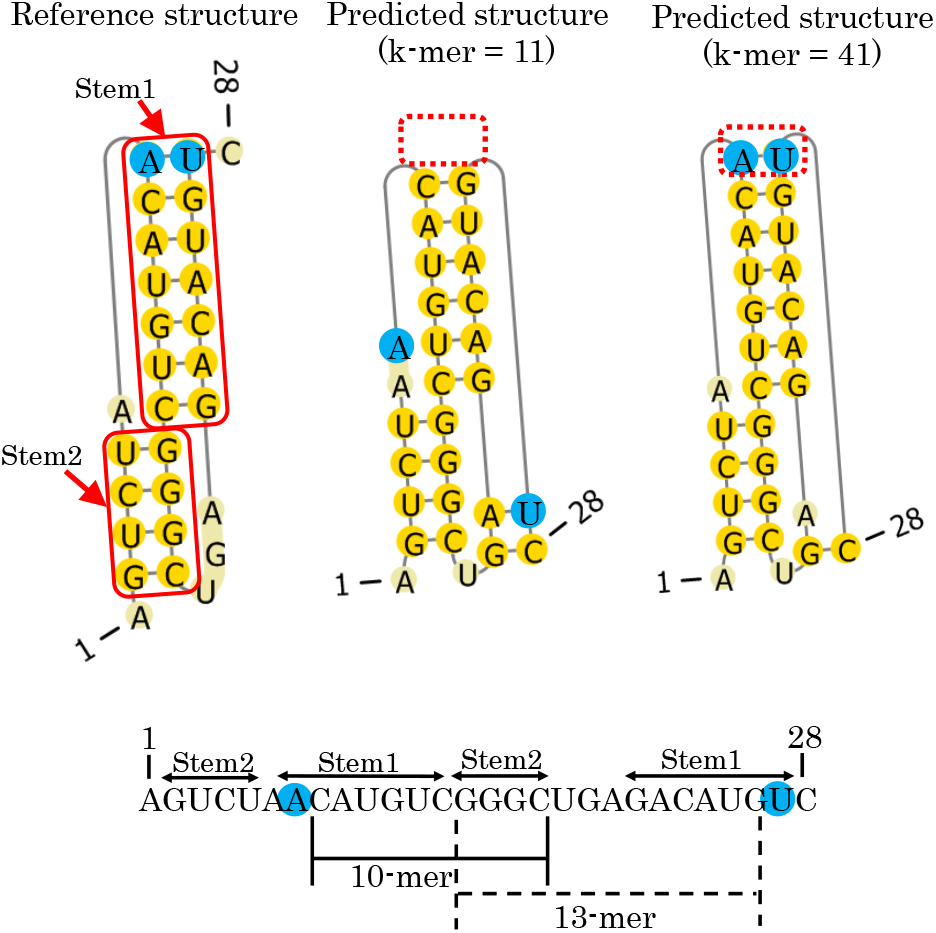
(Top) Comparison between the reference structure of ID : PKB189 (top-left) and predicted structures with the context length *k* = 11 (top-middle) and *k* = 41 (top-right). (Bottom) Distance between two stems (Stem 1 and Stem 2) in the pseudoknotted structure.

## Acknowledgements

The supercomputer system was provided by the National Institute of Genetics (NIG), Research Organization of Information and Systems (ROIS).

## Footnotes

1 https://www.ibm.com/analytics/data-science/prescriptive-analytics/cplex-optimizer

## References

Akutsu, T. (2000). Dynamic programming algorithms for rna secondary structure prediction with pseudoknots. Discrete Applied Mathematics, 104(1), 45 – 62.

Andronescu, M., Condon, A., Hoos, H. H., Mathews, D. H., and Murphy, K. P. (2007). Efficient parameter estimation for RNA secondary structure prediction. Bioinformatics, 23(13), 19–28.

Andronescu, M., Bereg, V., Hoos, H. H., and Condon, A. (2008). RNASTRAND: the RNA secondary structure and statistical analysis database. BMC Bioinformatics, 9, 340.

Andronescu, M., Condon, A., Hoos, H. H., Mathews, D. H., and Murphy, K. P. (2010a). Computational approaches for RNA energy parameter estimation. RNA, 16(12), 2304–2318.

Andronescu, M. S., Pop, C., and Condon, A. E. (2010b). Improved free energy parameters for RNA pseudoknotted secondary structure prediction. RNA, 16(1), 26–42.

Bellaousov, S. and Mathews, D. H. (2010). ProbKnot: fast prediction of RNA secondarystructure including pseudoknots. RNA, 16(10), 1870–1880.

Brierley, I., Pennell, S., and Gilbert, R. J. (2007). Viral RNA pseudoknots: versatile motifs in gene expression and replication. Nat. Rev. Microbiol., 5(8), 598–610.

Cao, S. and Chen, S. J. (2006). Predicting RNA pseudoknot folding thermodynamics. Nucleic Acids Res., 34(9), 2634–2652.

Carvalho, L. E. and Lawrence, C. E. (2008). Centroid estimation in discrete highdimensional spaces with applications in biology. Proc. Natl. Acad. Sci. U.S.A., 105(9), 3209–3214.

Chen, X., He, S. M., Bu, D., Zhang, F., Wang, Z., Chen, R., and Gao, W. (2008). FlexStem: improving predictions of RNA secondary structures with pseudoknots by reducing the search space. Bioinformatics, 24(18), 1994–2001.

Dirks, R. M. and Pierce, N. A. (2003). Apartitionfunction algorithmfornucleic acid secondary structure including pseudoknots. J Comput Chem, 24(13), 1664–1677.

Dirks, R. M. and Pierce, N. A. (2004). An algorithm for computing nucleic acid base-pairing probabilities including pseudoknots. J Comput Chem, 25(10), 1295–1304.

Do, C. B., Woods, D. A., and Batzoglou, S. (2006). CONTRAfold: RNA secondary structure prediction without physics-based models. Bioinformatics, 22(14), e90–98.

Do, C. B., Foo, C.-S., and Ng, A. Y. (2007). Efficient multiple hyperparameter learning for log-linear models. In J. C. Platt, D. Koller, Y. Singer, and S. T. Roweis, editors, NIPS, pages 377–384. Curran Associates, Inc.

Fechter, P., Rudinger-Thirion, J., Florentz, C., and Giege, R. (2001). Novel features in the tRNA-like world of plant viral RNAs. Cell. Mol. Life Sci., 58(11), 1547–1561.

Gardner, P. P., Daub, J., Tate, J., Moore, B. L., Osuch, I. H., Griffiths-Jones, S., Finn, R. D., Nawrocki, E. P., Kolbe, D. L., Eddy, S. R., and Bateman, A. (2011). Rfam: Wikipedia, clans and the “decimal” release. Nucleic Acids Res., 39(Database issue), D141–145.

Hamada, M., Kiryu, H., Sato, K., Mituyama, T., and Asai, K. (2009). Prediction of RNA secondary structure using generalized centroid estimators. Bioinformatics, 25(4), 465–473.

Hirose, T., Mishima, Y., and Tomari, Y. (2014). Elements and machinery of noncoding RNAs: toward their taxonomy. EMBO Rep., 15(5), 489–507.

Huang, X. and Ali, H. (2007). High sensitivity RNA pseudoknot prediction. Nucleic Acids Res., 35(2), 656–663.

Kato, Y., Sato, K., Hamada, M., Watanabe, Y., Asai, K., and Akutsu, T. (2010). RactIP: fast and accurate prediction of RNA-RNA interaction using integer programming. Bioinformatics, 26(18), i460–466.

Lorenz, R., Bernhart, S. H., Honer Zu Siederdissen, C., Tafer, H., Flamm, C., Stadler, P. F., and Hofacker, I. L. (2011). ViennaRNA Package 2.0. Algorithms Mol Biol, 6, 26.

Lyngsø, R. B. and Pedersen, C. N. (2000). RNA pseudoknot prediction in energy-based models. J. Comput. Biol., 7(3-4), 409–27.

McCaskill, J. S. (1990). The equilibrium partition function and base pair binding probabilities for RNA secondary structure. Biopolymers, 29(6-7), 1105–1119.

Morita, K., Saito, Y., Sato, K., Oka, K., Hotta, K., and Sakakibara, Y. (2009). Genome-wide searching with base-pairing kernel functions for noncoding RNAs: computational and expression analysis of snoRNA families in Caenorhabditis elegans. Nucleic Acids Res., 37(3), 999–1009.

Nussinov, R., Pieczenick, G., Griggs, J., and Kleitman, D. (1978). Algorithms for loop matching. SIAM J. Appl Math., 35, 68–82.

Reeder, J. and Giegerich, R. (2004). Design, implementation and evaluation of a practical pseudoknot folding algorithm based on thermodynamics. BMC Bioinformatics, 5, 104.

Ren, J., Rastegari, B., Condon, A., and Hoos, H. H. (2005). HotKnots: heuristic prediction of RNA secondary structures including pseudoknots. RNA, 11(10), 1494–1504.

Reuter, J. S. and Mathews, D. H. (2010). RNAstructure: softwarefor RNAsecondary structure prediction and analysis. BMC Bioinformatics, 11, 129.

Rivas, E. (2013). The four ingredients of single-sequence RNA secondary structure prediction. Aunifying perspective. RNA Biol, 10(7), 1185–1196.

Rivas, E. and Eddy, S. R. (1999). A dynamic programming algorithm for RNA structure prediction including pseudoknots. J. Mol. Biol., 285(5), 2053–2068.

Ruan, J., Stormo, G. D., and Zhang, W. (2004). An iterated loop matching approach to the prediction of RNA secondary structures with pseudoknots. Bioinformatics, 20(1), 58–66.

Sato, K., Mituyama, T., Asai, K., and Sakakibara, Y. (2008). Directed acyclic graph kernels for structural RNA analysis. BMC Bioinformatics, 9, 318.

Sato, K., Hamada, M., Asai, K., and Mituyama, T. (2009). CENTROIDFOLD: a web server for RNA secondary structure prediction. Nucleic Acids Res., 37(Web Server issue), W277–280.

Sato, K., Kato, Y., Hamada, M., Akutsu, T., and Asai, K. (2011). IPknot: fast and accurate prediction of RNA secondary structures with pseudoknots using integer programming. Bioinformatics, 27(13), 85–93.

Sato, K., Kato, Y., Akutsu, T., Asai, K., and Sakakibara, Y. (2012). DAFS: simultaneous aligning and folding of RNA sequences via dual decomposition. Bioinformatics, 28(24), 3218–3224.

Schroeder, S. J. and Turner, D. H. (2009). Optical melting measurements of nucleic acid thermodynamics. Meth. Enzymol., 468, 371–387.

Staple, D. W. and Butcher, S. E. (2005). Pseudoknots: RNA structures with diverse functions. PLoS Biol., 3(6), e213.

Tsochantaridis, I., Joachims, T., Hofmann, T., and Altun, Y. (2005). Large margin methods for structured and interdependent output variables. J. Mach. Learn. Res., 6, 1453–1484.

Turner, D. H. and Mathews, D. H. (2010). NNDB: the nearest neighbor parameter database for predicting stability of nucleic acid secondary structure. Nucleic Acids Res., 38(Database issue), D280–282.

van Batenburg, F. H., Gultyaev, A. P., and Pleij, C. W. (2001). PseudoBase: structural information on RNA pseudoknots. Nucleic Acids Res., 29(1), 194–195.

Zakov, S., Goldberg, Y., Elhadad, M., and Ziv-Ukelson, M. (2011). Rich parameterization improves RNA structure prediction. J. Comput. Biol., 18(11), 1525–1542.

Zuker, M. (1989). On finding all suboptimal foldings of an RNA molecule. Science, 244(4900), 48–52.

